# Disentangling protein metabolic costs in human cells and tissues

**DOI:** 10.1101/2023.06.05.543683

**Authors:** Mónica Chagoyen, Juan F Poyatos

## Abstract

Many intrinsic functional characteristics of cells and tissues shape their genome-wide expression patterns. But what other factors might also modulate these patterns are not fully known. Here, we revisit the general model of costs in which the protein products of highly expressed genes should be short and made up of biosynthetically cheap amino acids. We first use single-cell expression data from a collection of human cell types to confirm this model with a twist: the most highly expressed proteins tend to be particularly short and use expensive amino acids. By clustering how these two factors change with expression across all cell types, we identified a set of archetypal profiles that uniquely balance costs and that occur at different proportion across cell types. Similar profiles were also found by examining the expression data of tissues, which allowed us to recognize those following a more or less costly strategy. We then asked how this model might delineate the expression changes seen in a tumor relative to its normal solid tissue, as it has been argued that energy constraints determine cancer progression. We discovered a strong signal for the overexpression of biosynthetically cheap compact genes in cancer tissues. Our work highlights how both aspects of the metabolic cost of a protein, length and amino acid biosynthesis, represent valuable measures for understanding the different levels of biological organization and also the differences between health and disease.

## Introduction

It is challenging to define which properties determine the essential characteristics of a cell or tissue type (Trapnell 2015; Zeng 2022; Domcke and Shendure 2023). Traditionally, cells and tissues has been classified by functional or structural properties (Hall and Olson 2006; Vickaryous and Hall 2006). Later, this classification incorporated molecular characteristics with the idea that the different types could be catalogued based on the patterns of gene expression (Arendt 2008). Within this molecular framework, one could focus on the production and maintenance of the necessary proteome.

But the proteome composition can be influenced by several factors. On the one hand, there may be functional constraints that modulate it in a particular way (Lundberg et al. 2010). On the other hand, associated energy costs could also play a role since cellular or tissue resources are limited (Yang et al. 2021). This second aspect is not easy to estimate since there are many processes involved (Lynch and Marinov 2015), for example, the maintenance/stability of proteins, their translation costs, the biosynthesis costs of the constituent amino acids (AAs), etc.

We will consider here a unique feature of energetics, the energetic costs (ECs), or metabolic efficiency, of protein biosynthesis. In fact, it has previously been shown that this characteristic determines a bias in the abundance of AA in the proteome of unicellular organisms promoted by selection (Akashi and Gojobori 2002; Swire 2007). Likewise, this signal was also observed in multicellular organisms mediated by a high protein abundance despite the *a priori* view of a weak evolutionary force towards metabolic efficiency. The result was similarly reported in humans, but note that in this case some AAs are obtained through diet alone. Still, for the latter there is an opportunity cost that incorporates the incomplete use of precursors (Swire 2007; Heizer, Raymer, and Krane 2011). Note that highly expressed genes were argued to avoid long sequences to reduce translational costs. By controlling the length of the sequence, costs also depend on the specific AAs a protein is made of. But the bias appeared to be weak (Urrutia and Hurst 2003). However, more recently, a stronger anti-correlation between AA cost and abundance was reported, indicating that this effect is ultimately driven by autotroph biosynthetic cost (Zhang et al. 2018).

Beyond the precise explanation of metabolic efficiency in humans (which could be caused by selection, mutational constraints and/or some functional limitations as described above), we will use the EC of a protein as a potential metric to characterize distinguished classes of cells or tissues. Furthermore, the enhanced need for AAs to support cancer progression suggests that this measure could be significant in the context of reading tumor progression (Tsun and Possemato 2015). The view in this case is that the somatic evolution of cancer would not only minimize the expression of unwanted proteins but would also dictate an “economical” use of AAs (Zhang et al. 2018).

To evaluate the impact of the EC of a protein on genome-wide expression patters we will decouple it into what we termed two “compensation” schemes corresponding to the length and averaged AA biosynthetic costs (Fig. 1A, note that this second measure ultimately describes the utilization of the different AAs in the human proteome). Indeed, these two factors contribute to the EC of a protein, defined as the sum of the costs of each AA (*ci*) times its abundance (*ai*), i.e., 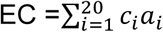. Since *ai*=*fiL*, with *fi* the relative abundance, frequency, of a particular AA, EC can be rewritten as EC = L*AEC. Here, L defines the first compensation strategy, and 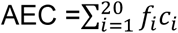 the average energy cost (AEC), the second.

**Figure 1.**
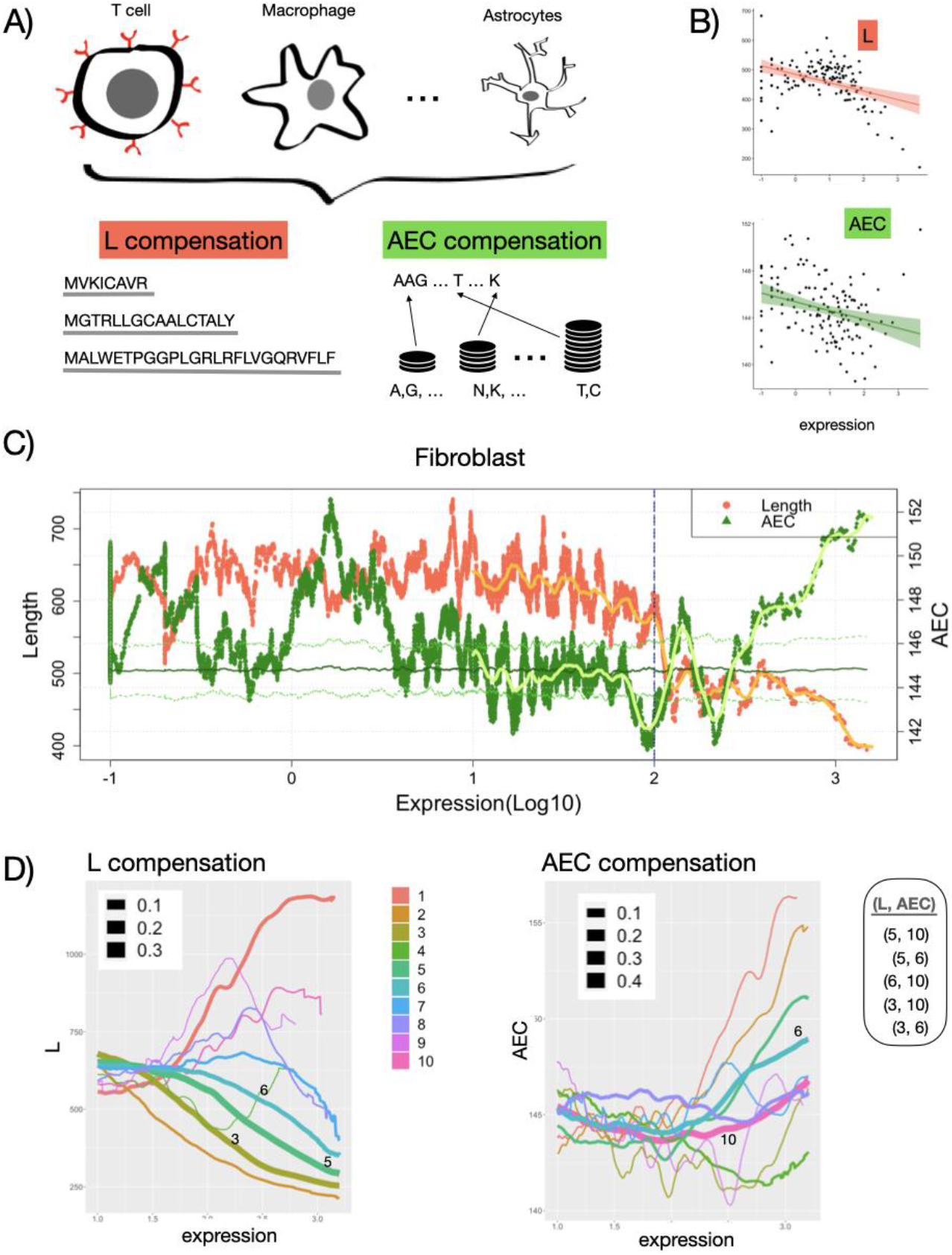
L- and AEC-based compensation in human cells. A) Different cell types balance the cost of protein production in different ways by using shorter peptides (L compensation) or cheaper constituent amino acids (average energy cost –AEC– compensation symbolized here as coins, see main text). B) A representative decay of L (number of amino acids) and AEC (ATP/time) with gene expression (log_10_, AU) contributing to a general pattern of reduced energy cost with high expression (added black regression line and shaded area indicating 95% confidence interval, t-cell data). C) Sliding window analysis of L (red) and AEC (green) for the exemplary case of fibroblasts with added kernel smoothing for log_10_ expression > 1 (solid lines). Note that in the high expression regime (>2) AEC increases beyond what would be expected by chance given the observed lengths in that regime (mean and +/-2 SD of null model in solid/dashed green lines). D) 10 archetypal patterns of L (left) and AEC (right) with expression (the width represents the relative abundance of each class). The 5 most common patterns (L, AEC) are also shown. Methods for details.

Our analysis verifies that highly expressed genes correlate with shorter proteins constituted by biosynthetically cheaper genes. However, this latter signal comes with a twist as we find that most highly expressed genes tend to exhibit expensive AAs. We can then identify archetypal profiles of how L and AEC changes with expression that help us characterize both cell types and tissues. By later focusing on which genes change expression in cancer, we identified that those that increase expression (compared to their matched normal tissue) are characteristically short and with low AEC, while large genes with high AEC generally decrease expression. More generally, transformed tissues avoid higher costs due to either higher AEC or Ls. In summary, the two main components of the energy cost of proteins, length and AA usage, help to identify defining characteristics of cells and tissues. We hope that our work reinforces interest in how various energy expenditures contribute to the distinct construction and preservation of cellular and tissue organization.

## Results

### Human cell types show archetypal patterns in the compensation of expression costs

We first established the relationship between L and AEC transcript expression levels in different cell types using data from the human protein atlas project (Karlsson et al. 2021). This data is collected in 444 cell *clusters* that are manually annotated as 78 distinct cell types. For example, there are 5 clusters annotated as dendritic cells, 22 clusters as macrophages, etc. (Methods). We calculated for each cluster how L and AEC changes with expression and typically found a tendency of both scores decreasing the higher the expression (Fig. 1B, also Fig. S1A), a result validating the metabolic efficiency model at the single-cell level.

However, for AEC, we noticed in many cases a reversal in this trend for high expression levels that could be captured with a sliding-window analysis (e.g., Fig. 1C). This trend is explained by an average increase in AEC at the strongest range of highly expressed levels together with the fact that some highly expressed genes exhibit idiosyncratic architectures (Ls very short, AECs very high) (Fig. S1B).

To evaluate these associations in a more systematic way, we grouped the patterns obtained through the analysis of sliding windows for each cell type and defined a set of 10 archetypal L and AEC classes (Methods). Five combinations of L and AEC profiles describe the majority of cell types (Fig.1D), with (L, AEC) = (5,10) being dominant. Overall, these profiles exhibit different degree of decay in the length and rise of biosynthetic cost for highly expressed genes and are presented differentially in each cell type (Fig.S2-S3 for the explicit distributions; these profiles can be summarized into two scores, Lav and AECav, Fig. S2B, to be discussed in the following sections). Note that there are particularly “expensive” combinations, such as (L, AEC) = {(5,6), (6,6)}, found in 16/40 fibroblasts, 13/46 t-cells, or 6/22 macrophages. These combinations are also found primarily in early and late spermatids and spermatocytes, which are already known to be “expensive” cells, what confirms our analysis.

### A subset of highly expressed genes shows an idiosyncratic pattern of high AEC and low L

We then asked which precise highly expressed genes determine the pattern (high AEC, low L). To this end, we identified for each cell type those highly expressed genes with high AEC and low L (Methods). This leads overall to a list of 46 *idiosyncratic* genes (Table S1) of which 13 are housekeeping genes (ATP6V0E1, COX6B1, UQCR11, UBL5, RPS27L, POLR2K, WDR83OS, MRPS21, RBX1, MRPL33, SPTSSA, SF3B5 and NDUFB3) and 6 are genes that “define” cell types, i.e., whose expression in one cell type is at least four-fold higher to that of all cell types (MT3, DEFB1, HAMP, SPRR2F, FAM24A, PROK1, Methods). Do cell-defining or housekeeping genes predominantly show a high AEC and low L pattern in general? We found that housekeeping genes are, on average, highly expressed genes (average over all cell types) with low AEC and low L (Eisenberg and Levanon 2013), while cell type-defining genes are, on average, low expression genes with high AEC and low L (Fig. 2).

**Figure 2.**
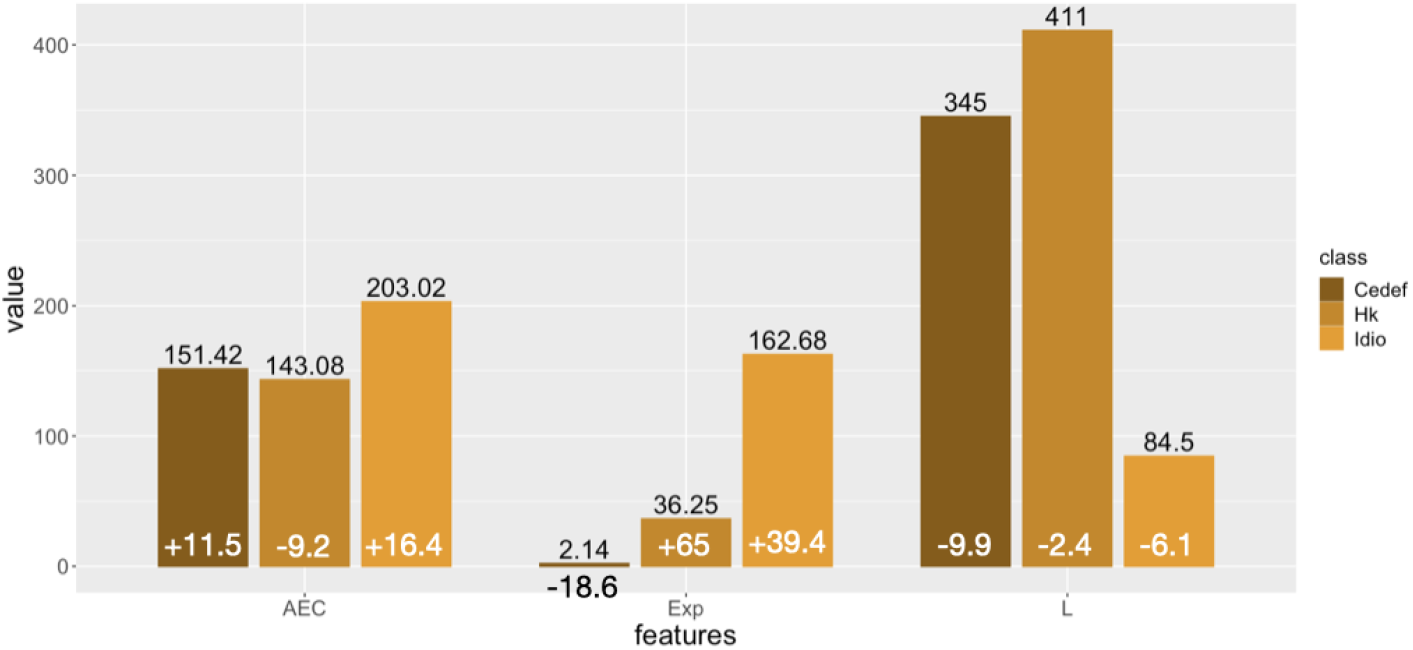
Median values of AEC, L, and mean expression for all cell types for the set of housekeeping (Hk), idiosyncratic (Idio) and cell-type defined (Cedef) genes. Inset (white) numbers indicate z values with respect to a random distribution obtain by 10000 permutations. Hk genes are cheaper, shorter and with higher mean expression in all cell types. Idio genes are by definition highly expressed, short, and expensive. Finally, cedef genes exhibit by definition less expression and are shorter and expensive. AEC, L and expression in ATP/time, number of AA and arbitrary units, respectively. Pairwise differences are all significant using the Wilcoxon signed-rank test. Number of each class: n_hk_ =3544, n_idio_=46, n_cedef_=2160.

### Human tissues show differences in the compensation of expression costs

Next, we consider to what extent these patterns are also observed in human tissues. Therefore, we now use human tissue expression data, bulk RNAseq data, to reanalyze the relationship between L, AEC, and transcript expression levels in 17,382 samples from 54 distinct GTEx tissues (GTEx Consortium 2017). Similar to human cell types, we found a trend of decreasing both scores the higher the expression. On the other hand, we also recovered the AEC signal increasing for high expression. To examine this systematically, we calculated L and AEC profiles with expression, as before, for each sample. We smoothed these profiles and pooled them to show 10 representative patterns (Fig. 3A). We notice in this case that two pairs of classes are dominant, Lclass-6 combined with AECclass-8 and Lclass-8 combined with AECclass-6 (Fig. S4).

**Figure 3.**
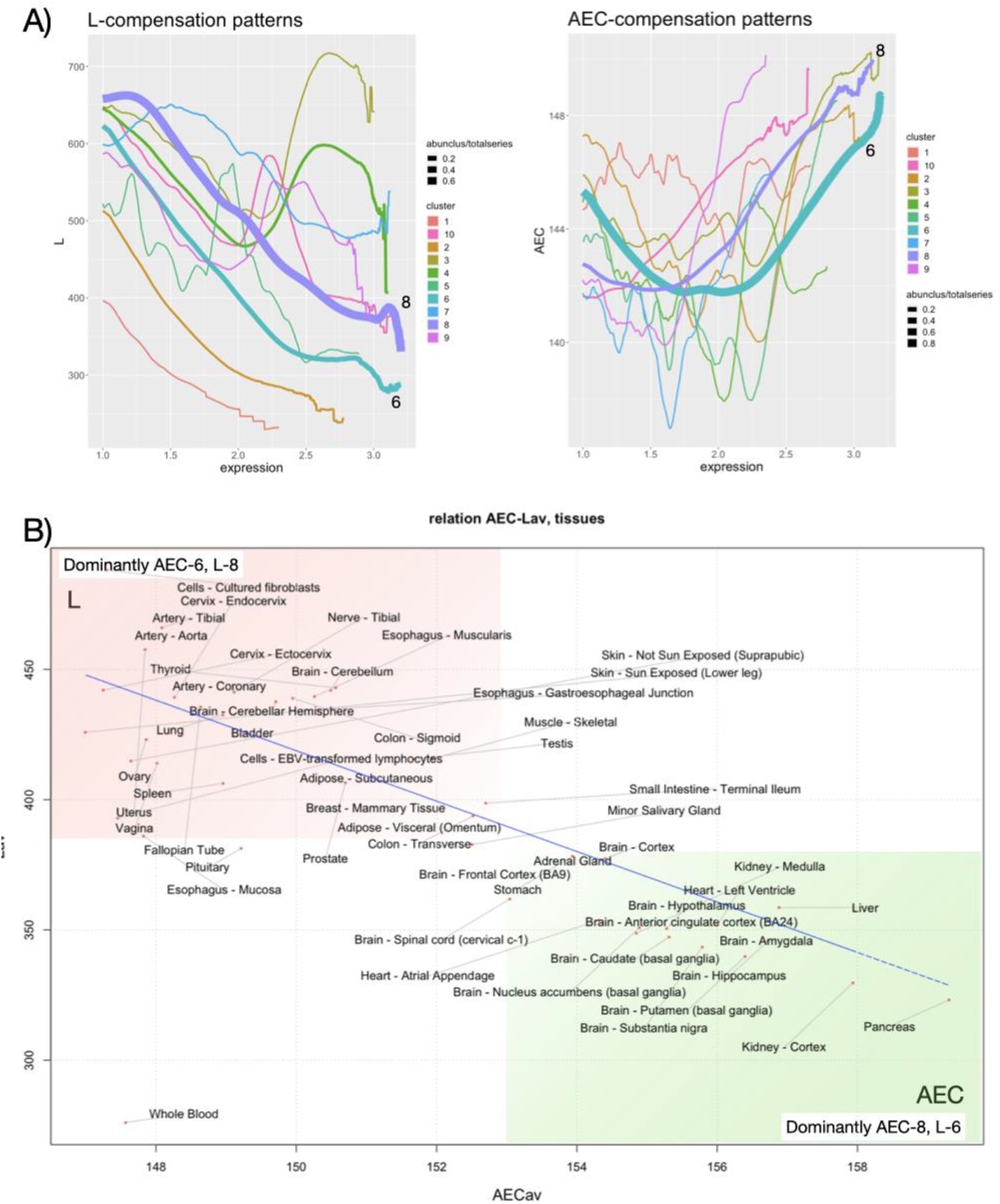
AEC- and L-based compensation in human tissues. A) Representative length decay and increase of AECs with expression (AU). B) AECav-Lav map of different human tissues. The bottom right corner locates tissues whose highly expressed genes are short but made up of metabolically expensive genes where the dominant pattern is (AECav, Lav) = (8,6). The upper left corner locates the opposite (highly expressed genes are long but made up of metabolically cheap genes) where the dominant pattern is (AECav, Lav) = (6,8). Red/green shading indicates a tendency to express long/expensive genes, respectively.

To understand these patterns and compare the different tissues more quantitatively, we consider two summary scores to describe each sample in the dataset, Lav and AECav, which indicate the weighted mean by (relative) expression of the L and AEC values for each gene, respectively. The amount of expression distributed among genes delimits the weights of the mean and therefore appreciably determines the value of the scores. When most genome expression is concentrated among highly expressed genes, this set is characteristically made up of compact genes with high AEC, leading to low Lav and high AECav (Fig. 3B), confirming the previously seen signal that highly expressed compact genes are also expensive (high AEC). Lav increases as high expression is distributed among a larger set of genes, with associated low AECav (Fig. 3B).

This representation thus identifies those tissues that use a higher compensation based on AEC or L and also which of them are more expensive overall. The latter is quantified by ECav, the average weighted by the (relative) expression of EC [this total cost is alternatively called ECcell in (Zhang et al. 2018)]. If we compare two tissues on the bottom right and top left of the map, for example, the pancreas *vs*. the uterus, we find that high expression is concentrated in a fewer number of genes in the pancreas than in the uterus (Fig. S5A, Methods). Thus, the pancreas seems to “compensate” the costs of high expression (highly expressed genes including, among others, pancreatic proteolytic enzymes) with short genes, while the uterus compensates it by using cheaper AA. Note that uterus total cost is stronger overall (higher ECav, Fig. S5B). Furthermore, the atypical nature of the blood is explained by the especially large contribution of the most expressed genes (Fig. S5). The four genes encoding human hemoglobin (HBB, HBA1, HBA2, HBD) account for an average of 42% of the total gene expression. They are significantly shorter (mean 144.5, p-value 2.9422e-04), but not significantly cheaper (AEC mean 146.42, p-value 0.1357).

Finally, we also focused on idiosyncratic highly expressed genes of low L and high AEC. These genes overlap considerably with those found before for the single-cell data (Fig. S6).

### Compensation of expression costs in cancer

Finally, an interesting context in which the two cost compensation strategies could become evident is that of cancer. Therefore, we obtained data for 16 normal paired samples from the TCGA portal (The Cancer Genome Atlas Research Network et al. 2013). Data analysis confirmed the above archetypal responses in L and AEC for all data (Fig. S7). We then ask to what extent tumor tissues differ from normal tissues in one or both strategies. To do this, we again use the AECav and Lav summary scores. Figure 4A shows an imagined distribution of a set of normal (solid) tissue samples and the possible directions in the (AECav, Lav) space in which it might change when the samples correspond to the tumor samples of the matched tissue (red arrows in Fig. 4A). We use this proposed space to classify data from the 16 TCGA pairs (Table S2).

**Figure 4.**
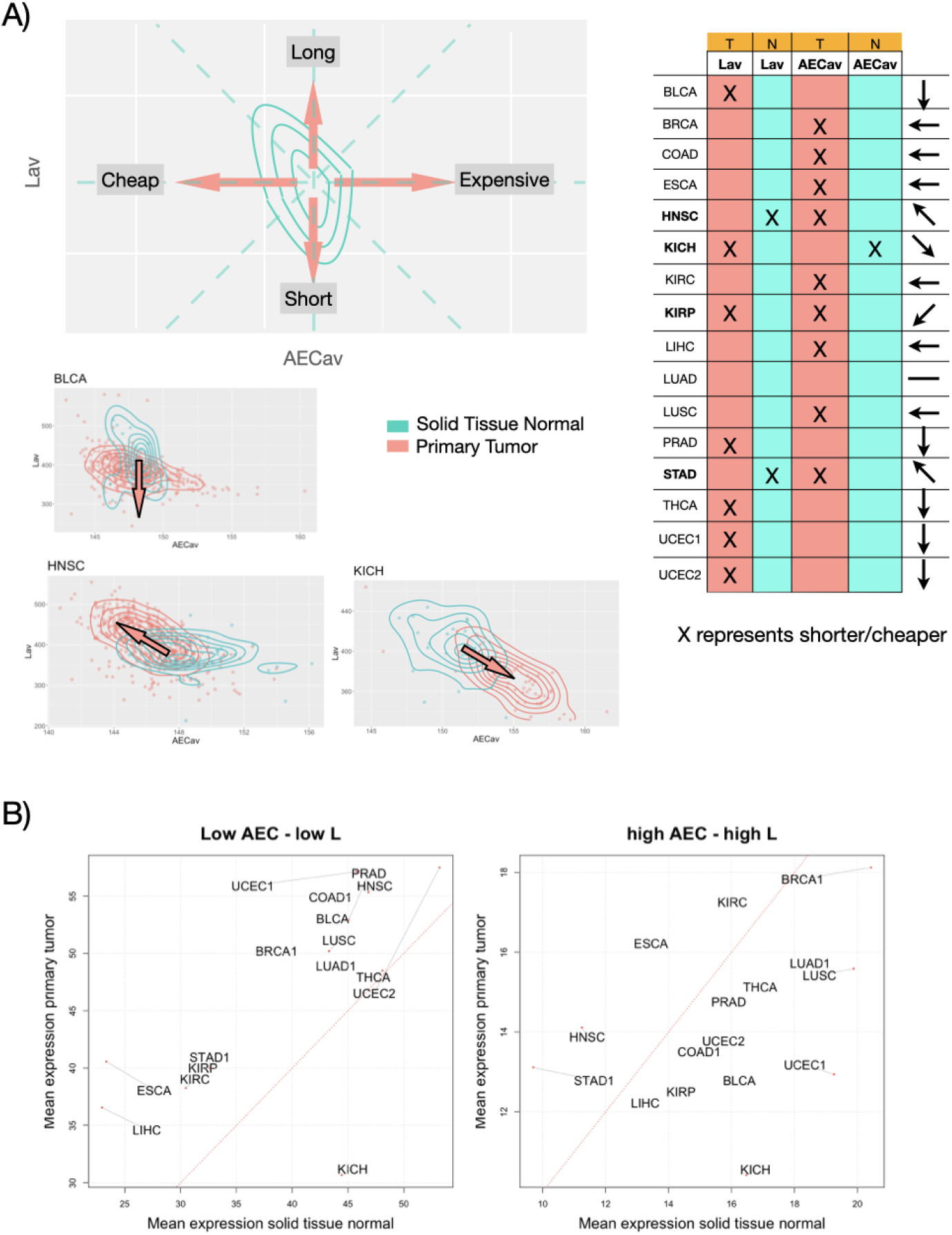
Tumors exhibit modified gene expression as a function of L and AEC. A) Top left. Cartoon of the possible shift of the distribution of a normal tissue sample in space (AECav, Lav) when compared to corresponding tumor samples of the same tissue. Bottom right. Three examples of this transition. Top right. Summary of the shift of tumor distribution to normal tissue for sixteen tumor types. The arrows in the right column indicated with respect to the map (AECav, Lav) on the left. B) Change in mean expression of AEC low - low L genes (left) or high AEC - high L genes (right) in tumors relative to normal tissues. The red line indicates slope = 1. In most cases, AEC low - L low genes increase their (mean) expression, while AEC high - L high genes decrease it.

We found that in 11 of 16 cases, tumor tissues save energy from highly expressed genes either by producing shorter proteins (5 cases) or by using cheaper AAs in proteins (6 cases). An extreme case is renal papillary cell carcinoma of the kidney (KIRP) in which both strategies are found (highly expressed genes are shorter and cheaper). Also notable are cases where there a savings in one strategy, but higher cost in the other (HNSC, KICH, STAD). Therefore, the overall picture is one of cost savings in tumor tissues, i.e., avoiding the upper right quadrant in Fig. 4A.

Instead of the sample distribution, we could also focus on how the “cheap” and “expensive” genes, in the AEC and L metrics, change their expression for each normal/tumor pair. We first divided genes into four categories that exhibit low or high (AEC, L) values, i.e., low-low, low-high, high-low, and high-high, respectively (categories selected here using 25/75% quantiles of the corresponding distributions, Methods). Then, for each normal/tumor pair, we obtained the averaged expression of the corresponding samples and calculated the mean of these averaged values for those genes within each AEC-L category. In particular, we found that almost all tumor samples (15 of 16) increased expression of “cheap” genes relative to their associated normal sample (AEC low – L low category, Fig. 4B left), while they decrease it (11 of 16) with respect to the “expensive” genes (high AEC – high L category, Fig. 4B right), which generalizes previous results (Zhang et al. 2018) to these two metrics. We also found that in nine cases tumor samples also upregulate genes that are low in AEC or L, even though they are costly in the contrasting score (low-high or high-low pairs).

## Discussion

Large amounts of genome-wide data now available with techniques such as single cell RNAseq or initiatives such as HPA or GTEX have reinforced interest in understanding the organization of living systems at the cell and tissue level. How can we condense all this data into knowledge? Our approach is to consider the energetic aspect of the proteome. This aspect has a long history which, in our discussion, is related to the studies that examined the biased use of AAs in the proteome (Akashi and Gojobori 2002; Swire 2007; Heizer, Raymer, and Krane 2011; Lynch and Marinov 2015; Zhang et al. 2018). These reports present a model in which abundant proteins are small and use biosynthetically cheap AAs. It is not entirely clear what forces contribute to this bias, although selection appears to be dominant in prokaryotes and, to a lesser extent, in eukaryotes, including humans.

We distangled here protein energy costs in two components: length (L) and average energetic cost of AAs (AEC). We have observed decreasing trends of L and AEC with higher expression in individual cell types and tissues, trends which were previously described using averaged and separate tissue data for L (Urrutia and Hurst 2003) and AEC (Zhang et al. 2018), respectively. Thus, these two cost-saving strategies appear to be general for most cell types, and not a composite property of tissues. Even so, we observed conflicting results for the L pattern in brain cells and tissues. While L increases with gene expression in cells (Fig S1), it generally decreases in tissues. This could be explained by the nature of the RNAseq data from brain cells (coming from single nuclei rather than from single cell for all other tissues). Alternatively, postmortem samples collected at GTEx could alter previous observations of increased expression of long genes in human and other metazoan brains (Ferreira et al. 2018; McCoy and Fire 2020).

We found archetypal patterns that appear more or less frequently in different cell types and that identify idiosyncratic genes that are short but made up of expensive AAs. As more expensive AAs tend to be highly conserved (Heizer, Raymer, and Krane 2011), this could imply that functional restrictions in this set of highly expressed genes suppose a burden to the cell than should be compensated by the rest of the proteome. Indeed, highly expressed proteins evolve slowly (Drummond et al. 2005).

Tissues also exhibit archetypal expression patterns and the use of two scores, AECav and Lav, helped us delineate which tissues balance energy cost using cheap AA or short proteins. Higher AECav is related to vital organs (liver, kidney, pancreas, heart and most brain tissues) with homeostatic and energy-related functions for the organism. These organs have also elevated metabolic rates (Wang et al. 2010). This can indicate that compensation of expensive AAs can also be performed at the organismal level. Finally, using paired samples of normal and cancer tissues, we also show that transformed tissues modulate costs using both AEC and L, generalizing previous reports (Zhang et al. 2018). That overexpressed genes in tumor samples are short and cheap, while long and expensive genes are often underexpressed, further corroborates this signal.

But although some of the previous results are consistent, their reading must necessarily be convoluted. Note that in the above we have used transcripts as an indicator of protein abundance. We expect general trends to be conserved at the proteome level as well, although this relationship is complex and protein abundance data obtained by mass spectroscopy are usually less precise (Liu, Beyer, and Aebersold 2016). Furthermore, we have only considered the cost of biosynthesis. Other protein costs are relevant: transcriptional initiation and elongation costs (the latter also depends on L); protein lifetimes are variable (Fornasiero and Savas 2023), etc.. The amount of AA protein that is recycled is not 100%, so new AA must be synthesized to maintain stable protein levels (Davies and Humphrey 1978; Smith et al. 1988) with distinct recycling levels in different tissues (leucine recycling 0.58 in whole brain; 0.47 in liver), etc.

Moreover, the very definition of a cell type is currently under review with the deluge of data coming from the RNAseq projects. In our case, we use manually defined cell types with the use of marker genes, cell types in which, for example, we ignore potential functional variability within each type (Zeng 2022). However, we found some cell types that exhibit less costly patterns than others, but also pattern diversity within the same type (Figs. S2-S3).

Still, the summary scores, Lav and AECav, helped us to identify different tissue classes, confirming our initial intuition that energy, or at least some aspects of it, matters. Regarding the cancer versus control analysis, we confirmed the relevance of energy in two complementary ways. Overall, our results stress the importance of studies on the energy burden of cells and tissues and on how different cost-saving strategies are eventually integrated at the organic level.

## Acknowledgements

This work was supported by grant PID2019-108096RB-C22 (MC) and PID2019-106116RB-I00 (JFP) from the Spanish Ministry of Economy and Competitiveness with European Regional Development Fund. The results shown here are part based upon data generated by the TCGA Research Network: https://www.cancer.gov/tcga.

## Data availability

jupyter notebooks (and corresponding datasets) available upon request (JFP).

## Methods

### Single-cell data and analysis

We downloaded transcript expression levels from the Human Protein Atlas (https://www.proteinatlas.org), which 78 cell types. Briefly, scRNA-seq data were processed using a clustering algorithm and each *cluster* was manually annotated based on >500 well-known tissue– and cell type–specific markers. All cells in a group were pooled and the average transcript per million was calculated for all protein-coding genes (Karlsson et al. 2021). To obtain the general model of energy cost reduction with large expression, for example, Fig. 1B, we divided genes into clusters of 100 based on expression levels to obtain Spearman’s correlation between median expression level and median AEC/L in each group (Fig. S1A). For sliding window analysis, we plot the mean expression and AEC/L scores for windows of genes of size 250. We kernel smoothed these patterns for log10 expression values > 1 (smoothing bandwidth = 0.1) to apply unweighted pair group method with arithmetic mean hierarchical clustering. The null model is calculated by obtaining AEC values from 100 randomizations of the constituent AAs following the distribution of the complete genome for a given length. We computed the Euclidean distance of the kernel smoothed L and AEC patterns to develop a UPGMA hierarchical clustering analysis fixing 10 groups (Fig. 1D).

### Protein sequences and costs

We obtained the protein sequences from the Ensembl *Homo sapiens* high coverage assembly GRCh38 (for the splicing variants we take the longest sequence). AAs costs are measured by the number of high-energy phosphate bonds required for synthesis. Costs of the 20 AAs of bacteria and yeast are highly correlated and also with the corresponding 11 nonessential human AAs (Zhang et al. 2018). We specifically used the decay rate-normalized biosynthetic costs for yeast (Krick et al. 2014; Zhang et al. 2018). The cost of the whole proteome (frequencies of each AA times costs) is 144.56 ATP/time (Fig. S1C).

### Housekeeping, cell-type defining and idiosyncratic genes

A list of human housekeeping genes was obtained from (Eisenberg and Levanon 2013). Cell type defining genes were obtained partially following (Karlsson et al. 2021). In short, a gene defines a cell type when its average expression in all groups of a given type is at least four times greater than the average expression in all other cell types. Finally, idiosyncratic genes are obtained by identifying highly expressed genes (log10 expression values > 2.5) whose length is less than the 2.3 quantile and AEC is greater than the 97.7 quantile.

### GTEx tissue data

We use RNA-Seq data from the Genotype-Tissue Expression (GTEX) project, a comprehensive dataset of tissue-specific gene expression (GTEx Consortium 2017). The full dataset includes 56200 transcripts measured in 17382 conditions. Archetypal L and AEC patterns obtained by hierarchical clustering as before.

### TCGA cancer data

We use RNA-seq data from the Cancer Genome Atlas Program (TCGA), a comprehensive resource of cancer tissue gene expression (https://portal.gdc.cancer.gov/) (The Cancer Genome Atlas Research Network et al. 2013). We analyzed only 16 cancer groups (15 cancer types) that contained both solid tumor and paired normal tissue data (Table S2). For the analysis of the expression change, we consider four classes of genes below or above the 25 and 75 quantiles, respectively as (low L, low AEC), (low L, high AEC), (high L, low AEC), and (high L, high AEC).

## Supplement

**Table S1**. List of idiosyncratic genes for single cells including their respective AEC, L, and mean expression (in all cell types).

**Table S2**. List of tumor-normal paired samples from TGCA. Number of samples normal (N) and tumor (T) also shown.

**Figure S1. Models of reduction of energetic costs with high expression**.

A) Spearman correlation coefficient of length *vs*. expression (x-axis) and AEC *vs*. expression (y-axis) after dividing genes into groups of 100 based on expression levels (Methods). Most of the clusters confirm a cost reduction model with large expression (negative correlations), either by reducing the length or using amino acids that are cheap to synthesize. For AEC, we only found 5 cases with positive correlations. For L, we found 57 cases of which 45 correspond to brain (all clusters related to brain). Highly expressed proteins with long sequences are here dominantly related to cell substrate junction organization. B) “AEC change” measures how much larger the (median) AEC is for the subset of the strongest highly expressed genes compared to the median of an equally sized random subset sampled from highly expressed genes (The -log_10_ p value of this nonparametric test is shown). “High AEC expression” quantifies the degree to which short (low L) and expensive (high AEC) genes are particularly highly expressed compared to the full set of low L genes (again, the measure gives the -log_10_ p value of this test). Dashed blue lines indicate p = 0.05. These measures show a general trend of an average increase in AEC at the strongest highly expressed levels together with the fact that the highly expressed genes are enriched for very short L and very high AEC. C) Cost-frequency relationship of the 20 amino acids.

**Figure S2. Properties of L and AEC characteristic patterns**. A) Distribution of cell clusters in the 10 different L and AEC characteristic patterns. See also Fig. 1D, main text. Lav and AECav scores (see Methods) summarizes the differences of the L and AEC patterns. Red filled circles denote the most abundant classes.

**Figure S3. Distribution of L and AEC classes in the different cell types**. Frequency of the different classes of L and AEC for each cell type together with the number of cell clusters (right curve).

**Figure S4. Distribution of L and AEC archetypal classes in the different tissue types**.

Frequency of the different classes (modes) of L and AEC for each tissue type.

**Figure S5. Lav – AEC map for tissues considering only those genes concentrating**

**∼50% of expression**. A) This map already shows a similar distribution to Fig. 3B and also highlights the percentage of genes (dot sizes) that carry ∼50% of the whole expression. The lower this percentage, the more concentrated the expression in a few genes and the lower Lav. B) This map shows the associated ECav score to A). Note that ECav ∼Lav.

**Figure S6. Idiosyncratic genes**. List of idiosyncratic genes (high AEC, low L) that appear in different cells and tissues. Value between 0 and 1 indicating the ratio of cells (or tissues) types in which they appear as idiosyncratic, i.e., RPL37A appears as idiosyncratic in most cells and tissues.

**Figure S7. L- and AEC-compensation patterns in TGCA samples**. Archetypal patterns of L and AEC change with expression (log 10) in all TGCA samples.

