## Supplement for "Disentangling protein metabolic costs in human cells and tissues"

[1] Computational Systems Biology group (CNB-CSIC), Madrid E-28049, Spain.

[2] Logic of Genomic Systems Laboratory (CNB-CSIC), Madrid E-28049, Spain.

### Table S1

Table S1 Idiosyncratic genes

| Name | Ensembl | AEC_Y20 (ATP/time) | L (AAs) | meanExp (AU) | Description of those gene appearing as idiosyncratic in all cell types |
|---|---|---|---|---|---|
| RPL41 | ENSG00000229117 | 217.0000 | 25 | 5474.382692 | This gene encodes a **ribosomal protein** that is a component of the 60S subunit. The protein, which shares sequence similarity with the yeast ribosomal protein YL41, belongs to the L41E family of ribosomal proteins. It is located in the cytoplasm. The protein can interact with the beta subunit of protein kinase CKII and can stimulate the phosphorylation of DNA topoisomerase II-alpha by CKII. |
| PRM1 | ENSG00000175646 | 211.0000 | 51 | 3958.908333 | |
| RPL39 | ENSG00000198918 | 194.4216 | 51 | 3195.167308 | This gene encodes a **ribosomal protein** that is a component of the 60S subunit. The protein belongs to the S39E family of ribosomal proteins. It is located in the cytoplasm. |
| UQCR11 | ENSG00000127540 | 195.3125 | 56 | 361.944231 | This gene encodes the smallest known component of the **ubiquinol-cytochrome c reductase complex**, which forms part of the mitochondrial respiratory chain. The encoded protein may function as a binding factor for the iron-sulfur protein in this complex. |
| MT1X | ENSG00000187193 | 340.6967 | 61 | 555.875000 | |
| MT1F | ENSG00000198417 | 340.4672 | 61 | 150.473718 | |
| MT1A | ENSG00000205362 | 348.5820 | 61 | 15.273718 | |
| MT1G | ENSG00000125144 | 333.4839 | 62 | 865.073718 | |
| DEFB4A | ENSG00000171711 | 194.5078 | 64 | 1.479487 | |
| POLR2K | ENSG00000147669 | 202.6940 | 67 | 111.740385 | |
| ATP8 | ENSG00000228253 | 211.2794 | 68 | 411.886538 | |
| DEFB1 | ENSG00000164825 | 209.4118 | 68 | 270.579487 | |
| MT1M | ENSG00000205364 | 223.4275 | 69 | 76.499359 | |
| MRPL33 | ENSG00000243147 | 211.9420 | 69 | 154.462179 | |
| SPTSSA | ENSG00000165389 | 203.2746 | 71 | 89.259615 | |
| SPRR2F | ENSG00000244094 | 228.5417 | 72 | 4.198718 | |
| UBL5 | ENSG00000198258 | 192.8699 | 73 | 440.636538 | This gene encodes a member of a group of proteins similar to **ubiquitin**. The encoded protein is not thought to degrade proteins like ubiquitin but to affect their function through being bound to target proteins by an isopeptide bond. The gene product has been studied as a link to predisposition to obesity based on its expression in Psammomys obesus, the fat sand rat, which is an animal model for obesity studies. |
| MT2A | ENSG00000125148 | 241.2329 | 73 | 1046.998718 | This gene is a member of the **metallothionein family of genes**. Proteins encoded by this gene family are low in molecular weight, are cysteine-rich, lack aromatic residues, and bind divalent heavy metal ions, altering the intracellular concentration of heavy metals in the cell. These proteins act as anti-oxidants, protect against hydroxyl free radicals, are important in homeostatic control of metal in the cell, and play a role in detoxification of heavy metals |
| CRIP1 | ENSG00000213145 | 223.1039 | 77 | 164.229487 | |
| CKS1B | ENSG00000173207 | 189.8291 | 79 | 58.588462 | |
| ATP6V0E1 | ENSG00000113732 | 208.8642 | 81 | 276.856410 | This gene encodes a component of **vacuolar ATPase** (V-ATPase), a multisubunit enzyme that mediates acidification of eukaryotic intracellular organelles. V-ATPase dependent organelle acidification is necessary for such intracellular processes as protein sorting, zymogen activation, receptor-mediated endocytosis, and synaptic vesicle proton gradient generation. V-ATPase is composed of a cytosolic V1 domain and a transmembrane V0 domain. |
| RPS27 | ENSG00000177954 | 188.7381 | 84 | 4149.568590 | |
| HAMP | ENSG00000105697 | 206.9881 | 84 | 21.896795 | |
| MT1H | ENSG00000205358 | 202.1059 | 85 | 197.296154 | |
| COX6B1 | ENSG00000126267 | 210.3779 | 86 | 525.312179 | Cytochrome c oxidase (COX), the terminal enzyme of the **mitochondrial respiratory chain**, catalyzes the electron transfer from reduced cytochrome c to oxygen. It is a heteromeric complex consisting of 3 catalytic subunits encoded by mitochondrial genes and multiple structural subunits encoded by nuclear genes. The mitochondrially-encoded subunits function in electron transfer, and the nuclear-encoded subunits may be involved in the regulation and assembly of the complex. |
| SF3B5 | ENSG00000169976 | 191.0756 | 86 | 160.748077 | |
| RPS29 | ENSG00000213741 | 213.0747 | 87 | 2146.953846 | This gene encodes a **ribosomal protein** that is a component of the 40S subunit and a member of the S14P family of ribosomal proteins. The protein, which contains a C2-C2 zinc finger-like domain that can bind to zinc, can enhance the tumor suppressor activity of Ras-related protein 1A (KREV1). It is located in the cytoplasm. Variable expression of this gene in colorectal cancers compared to adjacent normal tissues has been observed, although no correlation between the level of expression and the severity of the disease has been found |
| MRPS21 | ENSG00000266472 | 193.8621 | 87 | 181.669872 | |
| DEFB119 | ENSG00000180483 | 193.8920 | 88 | 8.795513 | |
| DYNLL1 | ENSG00000088986 | 190.5449 | 89 | 605.633333 | |
| CCL18 | ENSG00000275385 | 193.1966 | 89 | 6.816026 | |
| RPL37A | ENSG00000197756 | 195.8370 | 92 | 2498.741026 | This gene encodes a **ribosomal protein** that is a component of the 60S subunit. The protein belongs to the L37AE family of ribosomal proteins. It is located in the cytoplasm. The protein contains a C4-type zinc finger-like domain. |
| SCGB2A1 | ENSG00000124939 | 197.6263 | 95 | 62.139744 | |
| CCL20 | ENSG00000115009 | 197.3750 | 96 | 239.078846 | |
| BANF2 | ENSG00000125888 | 202.7577 | 97 | 20.038462 | |
| KRTAP3-1 | ENSG00000212901 | 246.1990 | 98 | 4.857692 | |
| NDUFB3 | ENSG00000119013 | 189.0357 | 98 | 161.130769 | |
| SCRG1 | ENSG00000164106 | 203.8622 | 98 | 12.570513 | |
| LELP1 | ENSG00000203784 | 244.9694 | 98 | 110.038462 | |
| RPS27L | ENSG00000185088 | 197.5850 | 100 | 333.024359 | This gene encodes a protein sharing 96% amino acid similarity with **ribosomal protein** S27, which suggests the encoded protein may be a component of the 40S ribosomal subunit. |
| IFI27L1 | ENSG00000165948 | 202.0571 | 105 | 28.272436 | |
| PROK1 | ENSG00000143125 | 202.1810 | 105 | 13.543590 | |
| FAM24A | ENSG00000203795 | 197.9000 | 105 | 1.583333 | |
| WDR83OS | ENSG00000105583 | 201.8349 | 106 | 189.899359 | |
| RBX1 | ENSG00000100387 | 223.1250 | 108 | 226.286538 | |
| MT3 | ENSG00000087250 | 224.9248 | 113 | 11.033333 | |

### Table S2

Table S2. Cancer Samples

| Study Abbreviation | Study Name | T | N |
|---|---|---|---|
| BLCA | Bladder Urothelial Carcinoma | 412 | 19 |
| BRCA | Breast invasive carcinoma | 1063 | 112 |
| COAD | Colon adenocarcinoma | 402 | 39 |
| ESCA | Esophageal carcinoma | 162 | 11 |
| HNSC | Head and Neck squamous cell carcinoma | 502 | 44 |
| KICH | Kidney Chromophobe | 65 | 25 |
| KIRC | Kidney renal clear cell carcinoma | 540 | 72 |
| KIRP | Kidney renal papillary cell carcinoma | 290 | 32 |
| LIHC | Liver hepatocellular carcinoma | 371 | 50 |
| LUAD | Lung adenocarcinoma | 499 | 54 |
| LUSC | Lung squamous cell carcinoma | 502 | 49 |
| PRAD | Prostate adenocarcinoma | 490 | 51 |
| STAD | Stomach adenocarcinoma | 343 | 30 |
| THCA | Thyroid carcinoma | 503 | 59 |
| UCEC1 | Uterine Corpus Endometrial Carcinoma | 407 | 19 |
| UCEC2 | Uterine Corpus Endometrial Carcinoma | 144 | 4 |

#### Figure S2

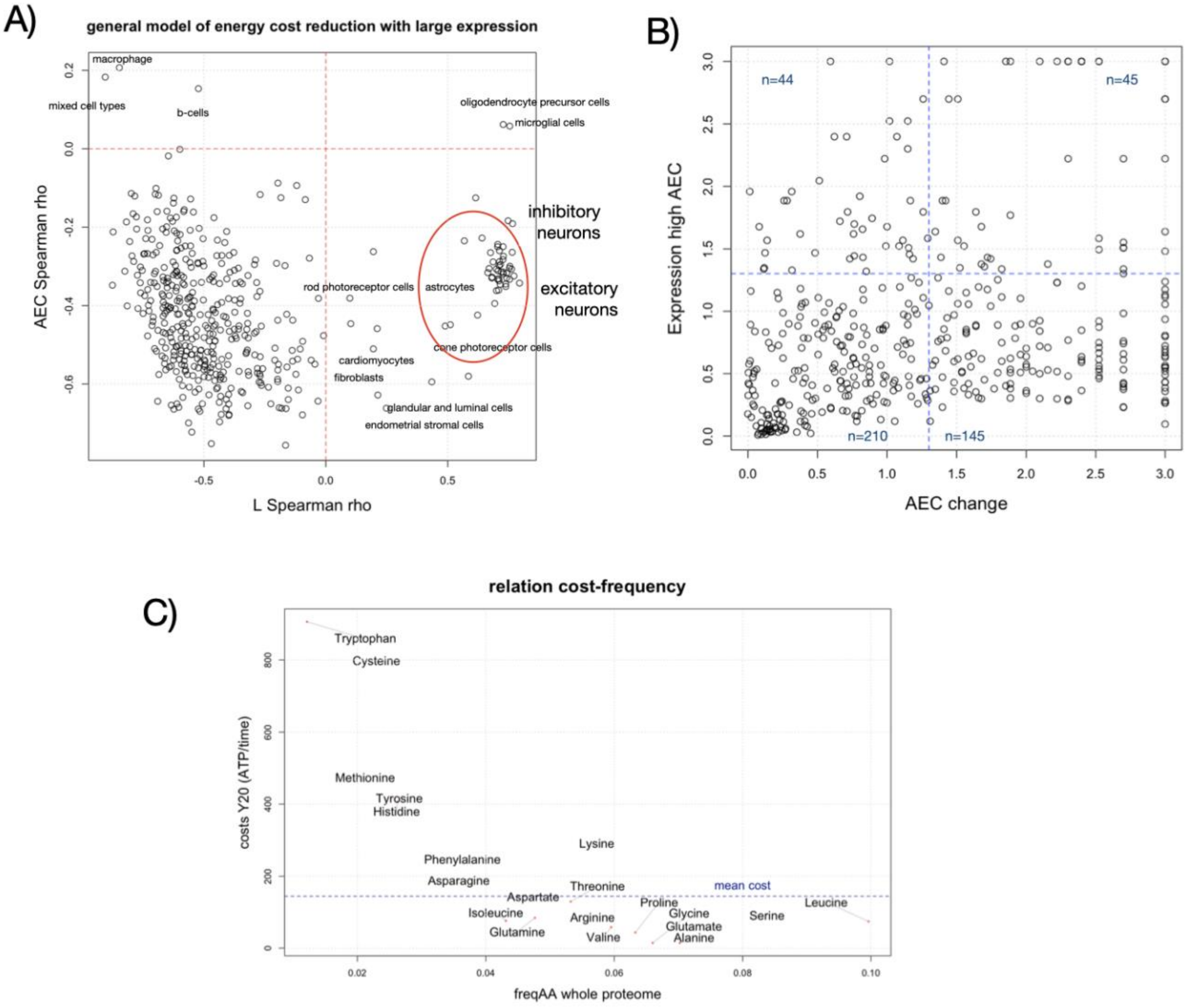

A)
general model of energy cost reduction with large expression
macrophage
mixed cell types
b-cells
oligodendrocyte precursor cells
microglial cells
inhibitory neurons
excitatory neurons
rod photoreceptor cells
astrocytes
cone photoreceptor cells
cardiomyocytes
fibroblasts
glandular and luminal cells
endometrial stromal cells
AEC Spearman rho
L Spearman rho
B)
n=44
n=45
n=210
n=145
Expression high AEC
AEC change
C)
relation cost-frequency
Tryptophan
Cysteine
Methionine
Tyrosine
Histidine
Lysine
Phenylalanine
Asparagine
Threonine
mean cost
Aspartate
Isoleucine
Glutamine
Arginine
Valine
Proline
Glycine
Glutamate
Alanine
Serine
Leucine
costs Y20 (ATP/time)
freqAA whole proteome

**Figure S3**

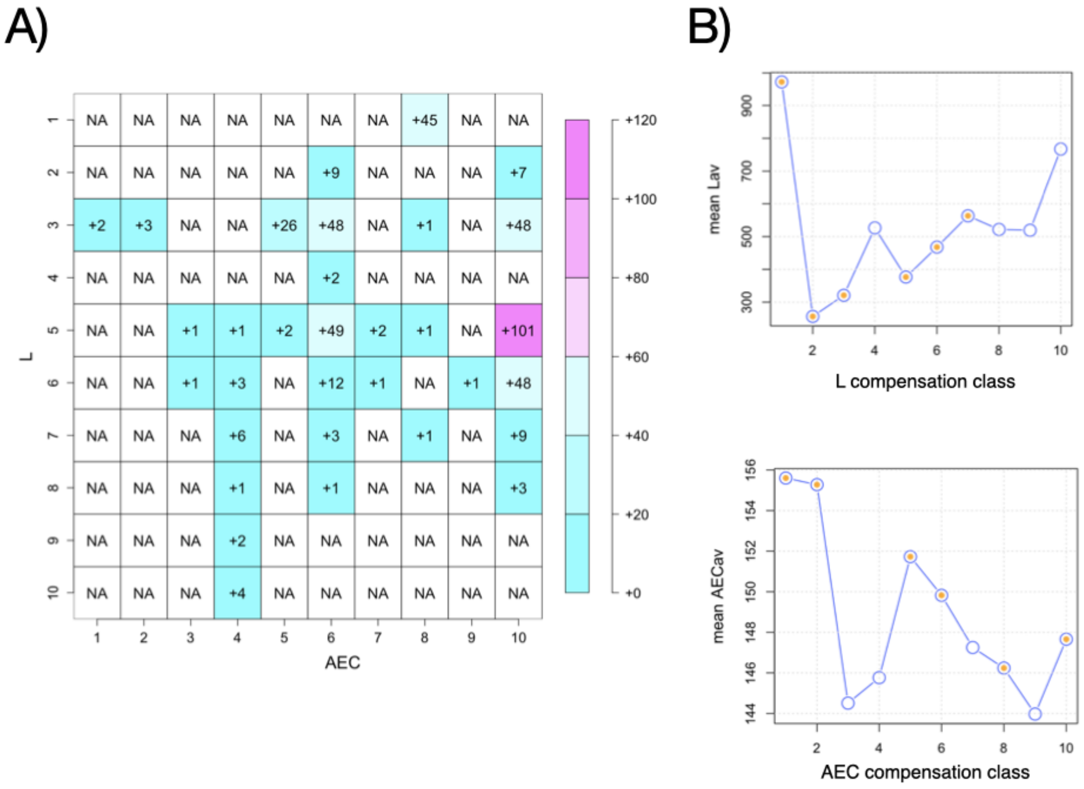
A)
NA NA NA NA NA NA NA +45 NA NA
NA NA NA NA NA +9 NA NA NA +7
+2 +3 NA NA +26 +48 NA +1 NA +48
NA NA NA NA NA +2 NA NA NA NA
NA NA +1 +1 +2 +49 +2 +1 NA +101
NA NA +1 +3 NA +12 +1 NA +1 +48
NA NA NA +6 NA +3 NA +1 NA +9
NA NA NA +1 NA +1 NA NA NA +3
NA NA NA +2 NA NA NA NA NA NA
NA NA NA +4 NA NA NA NA NA NA
L
AEC
+120
+100
+80
+60
+40
+20
+0
B)
mean Lav
L compensation class
mean AECav
AEC compensation class

#### Figure S4

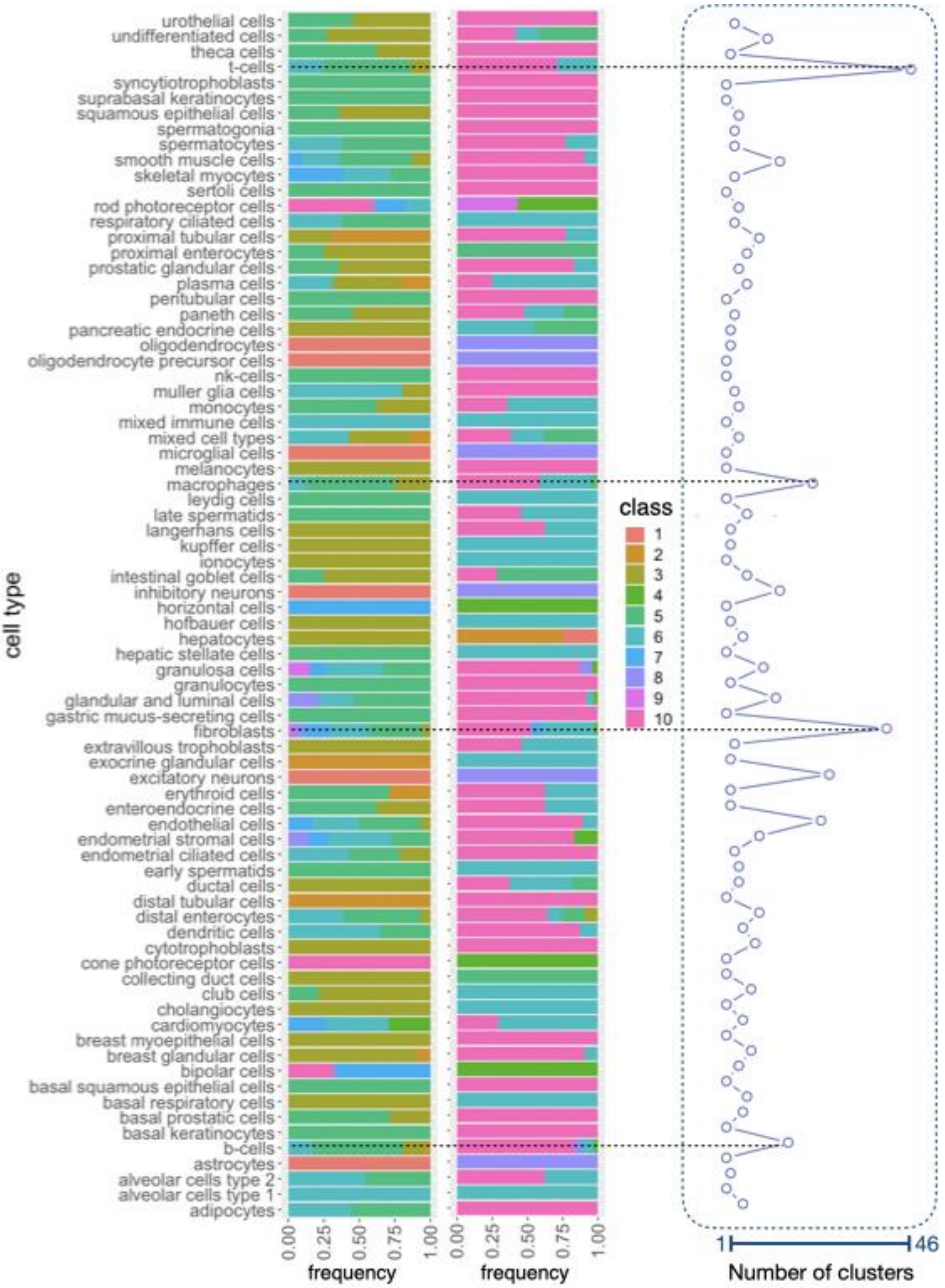

cell type
urothelial cells
undifferentiated cells
theca cells
t-cells
syncytiotrophoblasts
suprabasal keratinocytes
squamous epithelial cells
spermatogonia
spermatocytes
smooth muscle cells
skeletal myocytes
sertoli cells
rod photoreceptor cells
respiratory ciliated cells
proximal tubular cells
proximal enterocytes
prostatic glandular cells
plasma cells
peritubular cells
paneth cells
pancreatic endocrine cells
oligodendrocytes
oligodendrocyte precursor cells
nk-cells
muller glia cells
monocytes
mixed immune cells
mixed cell types
microglial cells
melanocytes
macrophages
leydig cells
late spermatids
langerhans cells
kupffer cells
ionocytes
intestinal goblet cells
inhibitory neurons
horizontal cells
hofbauer cells
hepatocytes
hepatic stellate cells
granulosa cells
granulocytes
glandular and luminal cells
gastric mucus-secreting cells
fibroblasts
extravillous trophoblasts
exocrine glandular cells
excitatory neurons
erythroid cells
enteroendocrine cells
endothelial cells
endometrial stromal cells
endometrial ciliated cells
early spermatids
ductal cells
distal tubular cells
distal enterocytes
dendritic cells
cytotrophoblasts
cone photoreceptor cells
collecting duct cells
club cells
cholangiocytes
cardiomyocytes
breast myoepithelial cells
breast glandular cells
bipolar cells
basal squamous epithelial cells
basal respiratory cells
basal prostatic cells
basal keratinocytes
b-cells
astrocytes
alveolar cells type 2
alveolar cells type 1
adipocytes
0.00
0.25
0.50
0.75
1.00
frequency
0.00
0.25
0.50
0.75
1.00
frequency
class
1
2
3
4
5
6
7
8
9
10
1
46
Number of clusters

### Figure S4

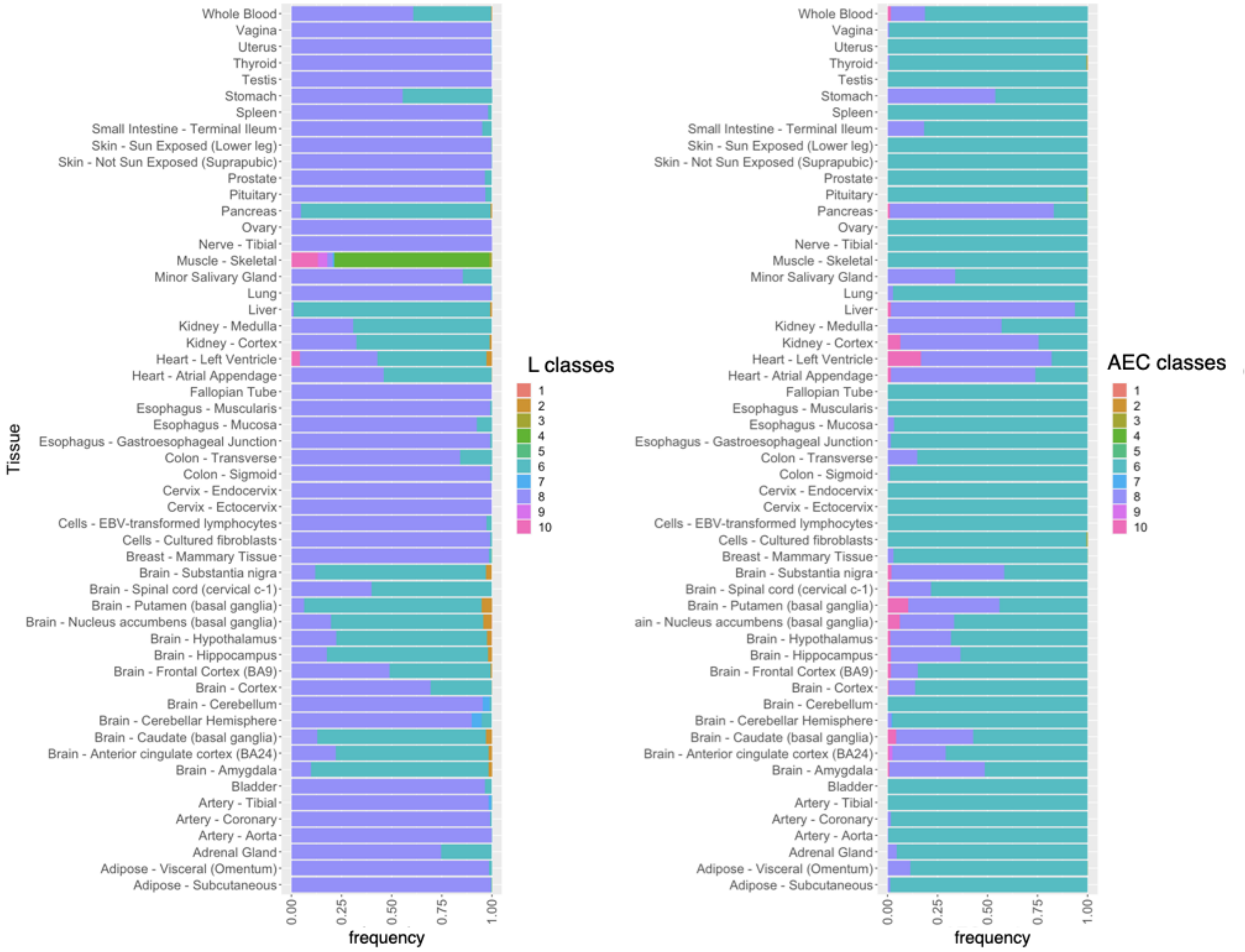

Whole Blood
Vagina
Uterus
Thyroid
Testis
Stomach
Spleen
Small Intestine - Terminal Ileum
Skin - Sun Exposed (Lower leg)
Skin - Not Sun Exposed (Suprapubic)
Prostate
Pituitary
Pancreas
Ovary
Nerve - Tibial
Muscle - Skeletal
Minor Salivary Gland
Lung
Liver
Kidney - Medulla
Kidney - Cortex
Heart - Left Ventricle
Heart - Atrial Appendage
Fallopian Tube
Esophagus - Muscularis
Esophagus - Mucosa
Esophagus - Gastroesophageal Junction
Colon - Transverse
Colon - Sigmoid
Cervix - Endocervix
Cervix - Ectocervix
Cells - EBV-transformed lymphocytes
Cells - Cultured fibroblasts
Breast - Mammary Tissue
Brain - Substantia nigra
Brain - Spinal cord (cervical c-1)
Brain - Putamen (basal ganglia)
Brain - Nucleus accumbens (basal ganglia)
Brain - Hypothalamus
Brain - Hippocampus
Brain - Frontal Cortex (BA9)
Brain - Cortex
Brain - Cerebellum
Brain - Cerebellar Hemisphere
Brain - Caudate (basal ganglia)
Brain - Anterior cingulate cortex (BA24)
Brain - Amygdala
Bladder
Artery - Tibial
Artery - Coronary
Artery - Aorta
Adrenal Gland
Adipose - Visceral (Omentum)
Adipose - Subcutaneous
Tissue
0.00
0.25
0.50
0.75
1.00
frequency
L classes
1
2
3
4
5
6
7
8
9
10
Whole Blood
Vagina
Uterus
Thyroid
Testis
Stomach
Spleen
Small Intestine - Terminal Ileum
Skin - Sun Exposed (Lower leg)
Skin - Not Sun Exposed (Suprapubic)
Prostate
Pituitary
Pancreas
Ovary
Nerve - Tibial
Muscle - Skeletal
Minor Salivary Gland
Lung
Liver
Kidney - Medulla
Kidney - Cortex
Heart - Left Ventricle
Heart - Atrial Appendage
Fallopian Tube
Esophagus - Muscularis
Esophagus - Mucosa
Esophagus - Gastroesophageal Junction
Colon - Transverse
Colon - Sigmoid
Cervix - Endocervix
Cervix - Ectocervix
Cells - EBV-transformed lymphocytes
Cells - Cultured fibroblasts
Breast - Mammary Tissue
Brain - Substantia nigra
Brain - Spinal cord (cervical c-1)
Brain - Putamen (basal ganglia)
ain - Nucleus accumbens (basal ganglia)
Brain - Hypothalamus
Brain - Hippocampus
Brain - Frontal Cortex (BA9)
Brain - Cortex
Brain - Cerebellum
Brain - Cerebellar Hemisphere
Brain - Caudate (basal ganglia)
Brain - Anterior cingulate cortex (BA24)
Brain - Amygdala
Bladder
Artery - Tibial
Artery - Coronary
Artery - Aorta
Adrenal Gland
Adipose - Visceral (Omentum)
Adipose - Subcutaneous
0.00
0.25
0.50
0.75
1.00
frequency
AEC classes
1
2
3
4
5
6
7
8
9
10

#### Figure S5

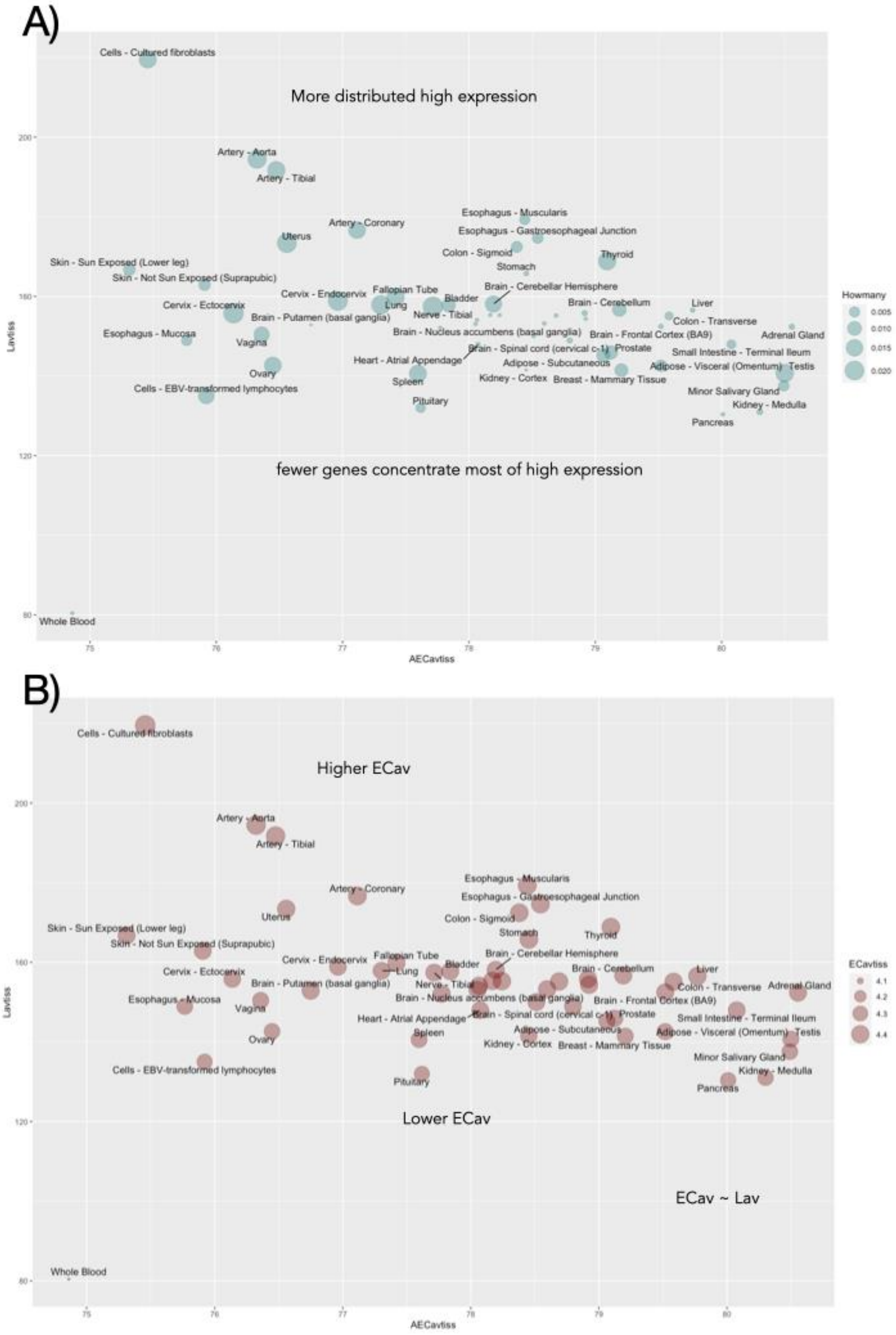

A)
Cells - Cultured fibroblasts
More distributed high expression
Artery - Aorta
Artery - Tibial
Artery - Coronary
Uterus
Esophagus - Muscularis
Esophagus - Gastroesophageal Junction
Colon - Sigmoid
Thyroid
Stomach
Skin - Sun Exposed (Lower leg)
Skin - Not Sun Exposed (Suprapubic)
Brain - Cerebellar Hemisphere
Cervix - Endocervix
Fallopian Tube
Bladder
Lung
Cervix - Ectocervix
Brain - Cerebellum
Liver
Nerve - Tibial
Brain - Putamen (basal ganglia)
Colon - Transverse
Esophagus - Mucosa
Brain - Nucleus accumbens (basal ganglia)
Brain - Frontal Cortex (BA9)
Adrenal Gland
Vagina
Brain - Spinal cord (cervical c-1)
Prostate
Small Intestine - Terminal Ileum
Heart - Atrial Appendage
Adipose - Subcutaneous
Adipose - Visceral (Omentum)
Testis
Ovary
Spleen
Kidney - Cortex
Breast - Mammary Tissue
Cells - EBV-transformed lymphocytes
Minor Salivary Gland
Pituitary
Kidney - Medulla
Pancreas
fewer genes concentrate most of high expression
Whole Blood
Lavtiss
AECavtiss
Howmany
0.005
0.010
0.015
0.020
B)
Higher ECav
Lower ECav
ECav ~ Lav
ECavtiss
4.1
4.2
4.3
4.4

#### Figure S6

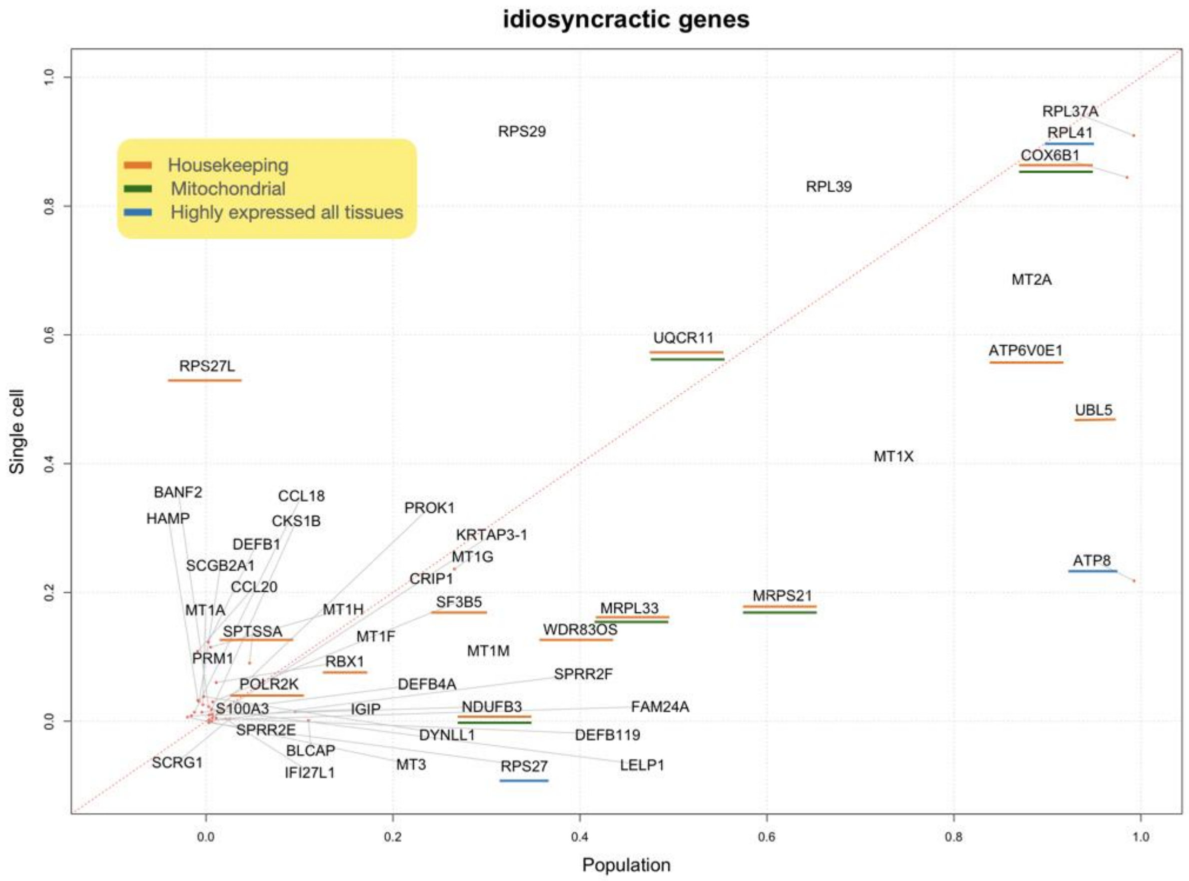


#### Figure S7

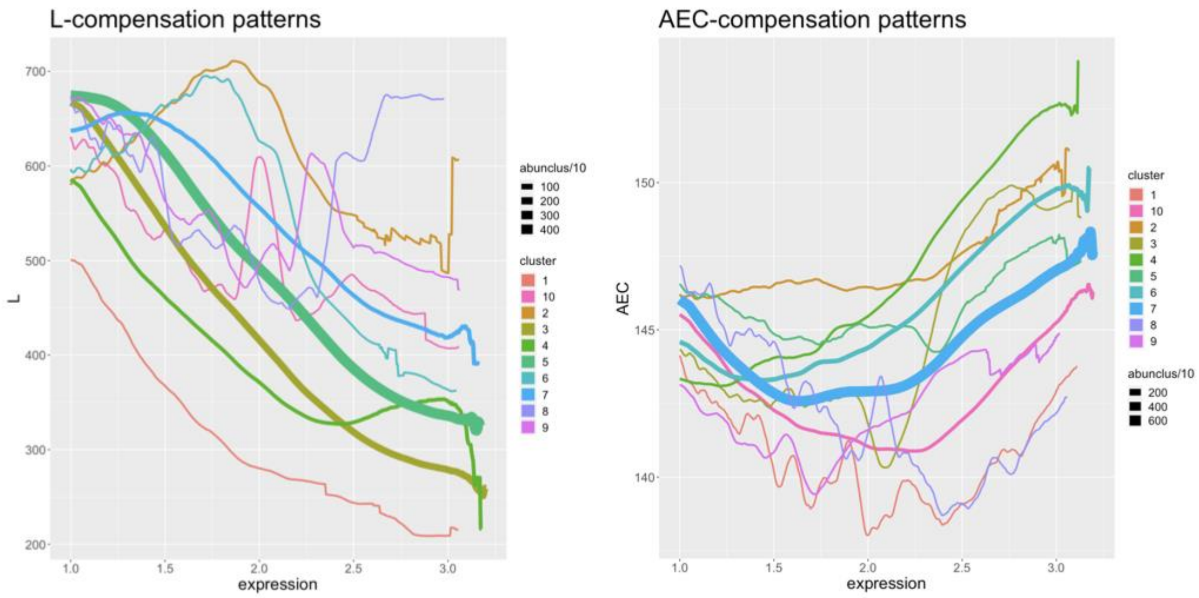
